# Neural correlates of the subjective experience of free-will during value-based risky decisions: a pilot study

**DOI:** 10.64898/2026.07.12.738094

**Authors:** Priyamvada Modak, Joshua W. Brown

## Abstract

Underlying the first-person notion of choice is the fundamental idea of ‘free-will’, which is challenged by decision-neuroscience aiming to explain and predict choices as outcomes of neural processes embedded within an ongoing chain of physical causes. The question of whether any choice is truly a free-choice and born out of free-will has long been a subject of philosophical debate. In this work, we do not take a position on this debate; rather we investigate the subjective experience of free-will, whose existence is more universally accepted. We had healthy participants report the level of their experienced free-will while performing a value-based risky decision-making task during an fMRI session to find the neural and behavioral correlates of this experience. We identified regions in mid-cingulum and middle frontal gyrus showing positive association with self-reported free-will as well as a region in hippocampus and parahippocampal gyrus showing a negative association. The requirement to report the experience of free-will was associated at a trend-level with a higher BOLD signal in striatum during decision-making. Behaviorally, we found a positive trend between RT and free-will. While our sample size is small, these results help form hypotheses for further studies with larger cohorts and provide a proof-of-concept for the investigations of neural and behavioral mechanisms of the subjective experience of free-will in decision research.

## INTRODUCTION

Whether we truly choose our actions and possess any free-will has garnered considerable yet unresolved philosophical debate (O’Connor and Franklin 2022). For the neuroscience approaches, aiming to explain volitional actions and behaviors as the product of interacting neurons (Hanes and Schall 1996), the debate is quite relevant. Consequently, the historical neuroscience focus on free-will has predominantly been to directly resolve this debate (Libet et al. 1983; Schlegel et al. 2015; Lavazza 2016), for example, via Libet-style experiments comparing the temporal profile of neural signals and volitional action.

While the nature and existence of free-will continues to be debated, the experience of it seems universal (Sarkissian et al. 2010; Robertson 2017). In this work, we did not take a position on the ontological status of free-will. We were interested not in whether free-will exists, but in the neural and behavioral correlates of the experience of it. The experience of free-will can be defined as the perceived capacity to choose freely among alternatives without constraints (Feldman 2017).

There is a substantial literature focused on neural mechanisms of free choice where the condition of participants making their own choice is contrasted with when they are instructed to select a specific option (Si, Rowe, and Zhang 2021; Thimm et al. 2012; Goldberg, Ullman, and Malach 2008; Welniarz et al. 2021; Walton, Devlin, and Rushworth 2004). This research is critical in informing our hypothesis for neural correlates of the experience of free-will but does not directly provide them. The neural correlates identified in these approaches correspond to perception, evaluation, and comparison of options which may not all be necessary for the experience of free-will. The approach also assumes that the former condition, of own choice, necessarily produces a greater experience of free choice for everyone, as compared to the condition of instructed choice. However, how and the extent to which the conditions differ in terms of experience of free choice is subjective and may change from one individual to another. Someone who considers the reward at stake to be too low may not feel obligated to follow the instructions in the control condition, rendering the assumption of lower free-will in that condition inaccurate. The neural mechanisms of free-choice, defined objectively as in the case of above, have been shown to not necessarily be same as the neural mechanisms of free-choice that is subjectively defined (Filevich et al. 2013).

There is limited previous neuroscientific work on the subjective experience of free-will. Filevich et al. (2013) had subjects perform a random number generation task and provide a rating for experienced free-will afterwards, using which the experimenters performed a median-split of the trials into free-choice and instructed. They showed that a medial post-central region corresponds to the differences between conditions, thus defined. Rens, Bode, and Cunnington (2018) also investigated subjective experience of free-will demonstrating that a greater representation of option-availability in right middle and inferior frontal gyri was associated with higher rating for experienced free-will in their decision-task.

In this work, we investigated the neural correlates of subjective experience of free-will in value-based risky decision-making. We focused on the candidate sources for the subjective experience of free-will by identifying brain regions whose decision-related BOLD activity varied with the self-reported experience of free-will during decisions across participants. Participants performed a two-alternative forced-choice task in which they made their own choices between risky options in the task condition, while they were instructed to choose a specific option in the control condition. Afterwards, they rated their experienced free-will in both the conditions on a five-point scale. To motivate accurate self-report, there was no time pressure to provide these ratings, and our presentation of the scale was designed to discourage ratings based solely on conserving time or effort when providing the self-report.

## METHODS

### Participants

We recruited 12 healthy participants (Mean age = 23.3 years; 6 females; 6 Caucasian, 5 Asian, and 1 Other) via flyering around the Indiana University campus, to perform a risky decision-making task. These participants were part of a bigger cohort (n = 30) described in Modak and Brown (2026) and were included in other analyses reported in that paper.

### Task

The task (Figure 1) consisted of selecting between two options, a gamble and a sure thing. The gamble had two or three possible outcomes, with the outcome values and their probabilities shown to the participants on the screen. The sure thing had one outcome value, also shown to the participants – if the sure thing option was selected, they received this outcome with complete certainty. The outcomes values associated with both options were in terms of points. The side on which the gamble appeared, the orientation of the gamble stimulus, and the color of gamble and sure-thing stimuli were selected randomly for each trial.

**Figure 1.**
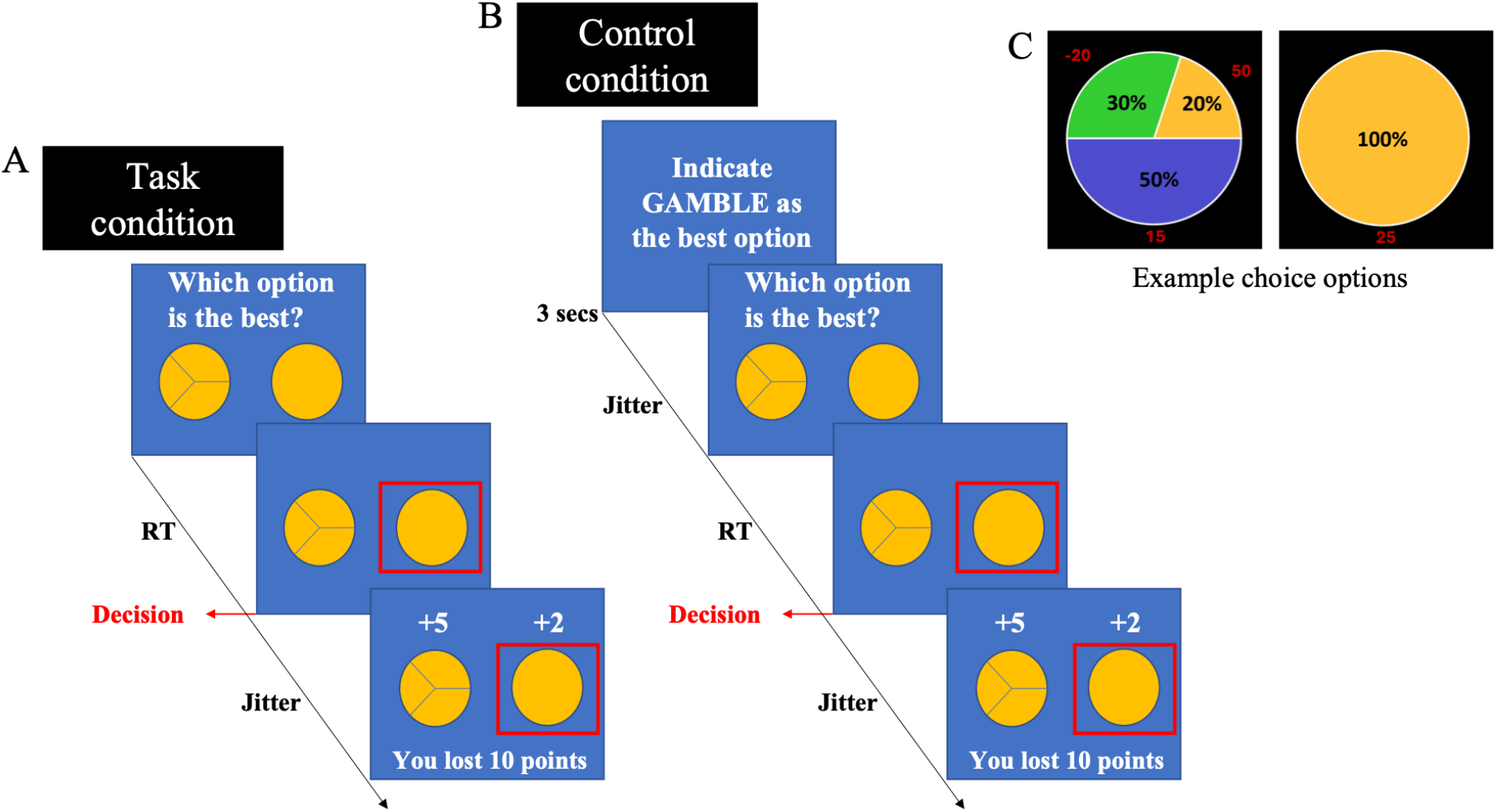
Progression of a trial in task condition (A) and control condition (B), where in every trial, a participant selected between a gamble and a sure thing (C). The gamble option entailed various possible outcomes, each with a corresponding probability accurately depicted on a pie chart.

In the task, as shown in figure 1A, participants were asked to indicate which option is the best, in the sense that it will actually pay more. The outcomes of both the options were shown to them in the feedback phase, and the points they received depended on how the outcome of the option they selected compared to the outcome of the option they did not select – they received 10 points if the outcome of selected option was more than or equal to the unselected option and lost 10 points otherwise. Thus, participants could maximize winnings by always choosing the option with a higher probability of a larger outcome value. This was the task condition, which we also refer to as the ‘Best’ condition. In the control condition, the participants selected between the same stimuli as the task condition, but before presenting the stimuli, participants were shown an instruction to select a specific option in the upcoming trial. They received 10 points if they selected an option as instructed and lost 10 points otherwise.

The control block was always conducted after the task blocks as the trials presented in this block were sampled from the trials presented in the task blocks. The progression of a control trial is shown in figure 1B. For any trial in the control block, participants were instructed to select the same option they had selected for that trial in the task block – they did not have to remember this nor were they explicitly asked to do so – but we made sure to instruct them to select the same option as was selected in the task block. This was done to minimize any confounds between the conditions that could arise from different stimuli presented or different probabilities of selecting a particular option or different response-stimuli combinations.

Participants had 5 seconds to respond in each case and were paid in dollars at the end proportional to points accumulated in each condition. More details on payment can be found in supplementary material (section 3). Each participant completed 2 blocks of the task condition followed by 1 block of control.

There were five different gamble stimuli and two different sure-thing values associated with it. Prior to the scan session, participants performed a certainty equivalent task online, which was used to identify the two sure-thing values for each gamble. The purpose of this task was to make sure that sure-thing values were such that the decisions weren’t too easy for a participant (supplementary material, section 2).

In any given trial, the gamble that was presented was chosen randomly from among the five possible gambles (supplementary material, section 1). The sure-thing was also chosen randomly from among the two sure-thing values associated with the gamble, as determined by the online certainty equivalent task, and gaussian noise was added to it.

### Free-will ratings

During the fMRI session, we asked participants to provide a rating on a scale of 1-5 for their experienced level of ‘free-will’ during the decisions. This was obtained at the end of every block. Please see figure 2 for the screens shown to the participants to obtain this rating. Participants were encouraged to think about internal and external constraints they had experienced while making their decisions and whether their choices could have gone the other way.

**Figure 2.**
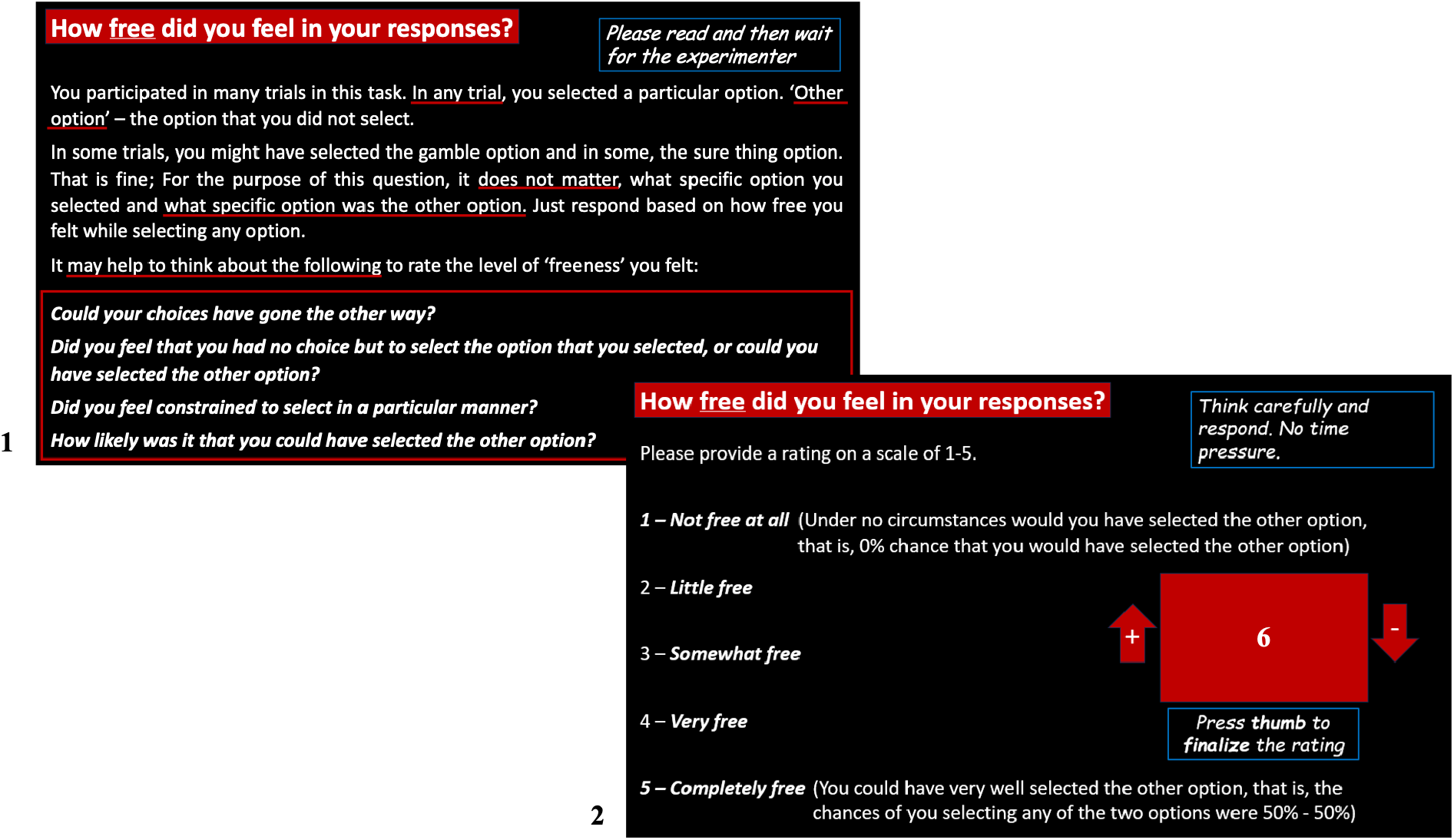
Screens shown to the participants to obtain free-will rating at the end of every block. Screen 1 was shown to the participants for roughly 60 seconds and then they were asked if they had read and understood everything and wanted to move to the next screen. They provided the rating on the next screen.

The initial rating on the screen was taken from the set {0, 2.5, 3.5, 6}. If participants submitted the initial rating, it was returned as ‘invalid’ and was not accepted, as the rating had to be a whole number between 1 and 5. The purpose of having an ‘invalid’ initial rating was to facilitate active thought about the rating, rather than passive acceptance of what was already on the screen just to move forward in the experiment. Participants had to add to or subtract from the initial rating using their index and middle fingers, respectively, to reach their intended rating. The mapping between finger (index/middle) and action (adding/subtracting) was randomly selected for each block. The initial rating was also randomly selected from the above set for each block, so that no rating was consistently ‘favored’ in terms of effort, that is, the number of button presses required to reach a valid rating.

At the end of the fMRI session. We also obtained a rating on the difficulty level of decision-making process for each task condition, on a scale of 1 – 10. Here, 1 was ‘Did not even feel like a decision; Low effort’, and 10 was ‘Very involved decision; High effort’.

### Behavioral analyses

We analyzed choice data to estimate the probability of choosing the sure-thing option for each participant in the task condition. The conflict between the options was quantified as, *p* ∗ (1 − *p*), where ‘p’ was the probability of choosing the sure-thing. We also quantified consistency in choice strategy. This is described in depth in Modak and Brown (2026). Briefly, we considered several candidate task strategies and performed model comparison to find the strategy that best described the behavior. This was performed at the level of individual subjects, so that the best fitting strategy could be different for different subjects. The ‘consistency’ for a particular subject was the extent to which the best fitting strategy accounted for the behavior and was quantified as the log-likelihood of the choice data given the strategy. A higher consistency, as quantified above, would indicate that a single strategy was consistently employed across the task while a smaller consistency would mean switching between multiple strategies or noisy behavior.

### fMRI acquisition and preprocessing

The fMRI data was collected using the Echo Planar Imaging (EPI) in the Siemens 3 Tesla TIM Trio MRI scanner in the Imaging Research Facility at Department of Psychological and Brain Sciences at Indiana University Bloomington. The data was obtained as axial slices at an angle on 30^°^ with anterior-commissure-posterior-commissure line using 64-channel head coil. In a single run lasting 6 mins, 178 volumes of the T2* weighted functional scans were collected with TR = 2000 ms, TE = 25 ms, flip angle = 70^°^, and 64 x 64 voxel matrix. A single volume consisted of 35 slices of thickness 3.8 mm.

One T1-weighted structural scan was collected with TR = 1800 ms, TE = 2.7 ms, flip angle = 9°, and 256 x 256 voxel matrix. This consisted of 160 slices of thickness 1 mm. Preprocessing was performed in SPM12. AFNI’s 3dDespike was used for spike correction. The scans were normalized to the standard Montreal Neurological Institute (MNI) space. Spatial smoothing was performed using 8 mm3 full-width-at-half-maximum (FWHM) kernel.

Due to a technical issue, we did not have the scanner start time for four participants. We used a machine learning approach to predict the scanner start time for the participants for whom this information was unavailable (Modak and Brown 2026). Briefly, a support vector machine was trained on fMRI data from the participants (n = 8) for whom we had the scanner start time and used it to predict the start time for other participants. The cross-validation prediction accuracy of this approach was 79%.

### fMRI analyses

General linear modeling (GLM) of fMRI data was performed using SPM12 in MATLAB (version 2013a). We created a GLM with ‘Decision’ as an event regressor at the time when participants submitted their decision. This was performed for the larger cohort (n = 30). Our main contrast of interest was (Decision_Task_ – Decision_Control_). We conducted whole-brain correlational analysis for the participants reported in this work (n = 12) to find associations between the main contrast and self-reported experience of free-will. We also performed small volume corrections of the above correlational analysis with the clusters showing significant effect of the main contrast in GLM analyses with the larger cohort.

We also performed two-sample t-tests of the main contrast values in the clusters above for the participants who were asked to report free-will ratings (n = 12) and for those from whom these ratings were not obtained (n = 18). Note that this paper reports on the free-will ratings and related neuroimaging and behavioral analysis and hence primarily focuses on the former group.

In addition to the task condition described above, which we call ‘Best’, participants also completed another task condition while in the scanner, referred to as ‘Choice’, in which they simply chose the option they preferred rather than the option most likely to yield a larger outcome. We defer treatment of this condition to Modak and Brown (2026), along with details of the whole cohort (n = 30) and the related behavioral and neuroimaging analyses.

## RESULTS

### Behavior

Self-reported free-will ratings in task condition were significantly greater than the control (Task > Control, mean = 1.12 units, non-parametric paired test, p = 0.041, n = 12). The individual variability in the extent to which these ratings differed between the conditions is shown in figure 3A. The ratings provided by each subject are also noted in the supplementary table S2. RT was longer (Task > Control, mean difference = 1500 ms, non-parametric paired test, p < 0.001, n = 12) and self-reported difficulty was higher (Task > Control, mean = 5.83 units, non-parametric paired test, p < 0.001, n = 12) in the Task vs. Control conditions.

**Figure 3.**
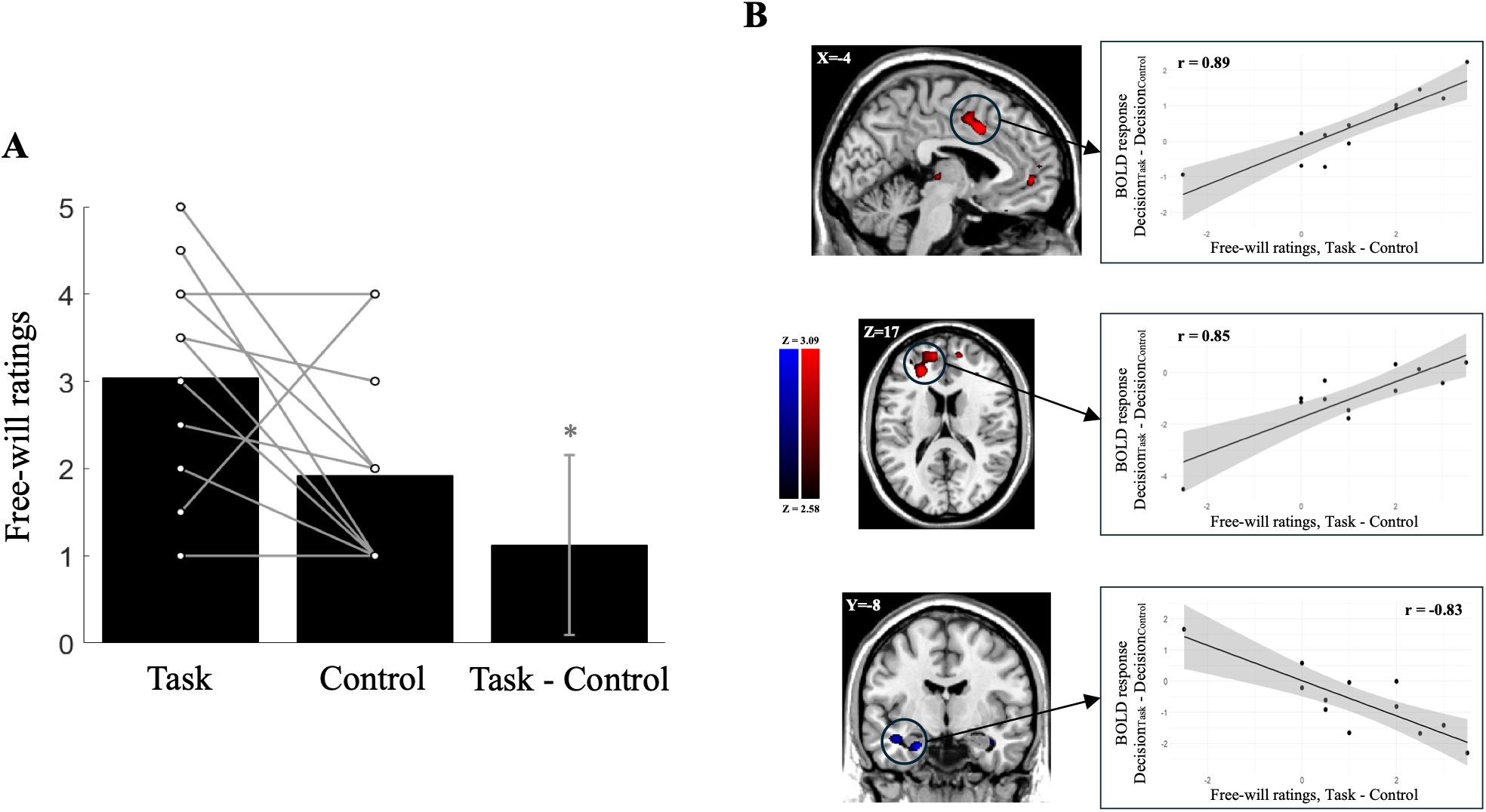
Neural correlates of experienced free-will at the time of decisions. A – Self-reported retrospective ratings of experienced free-will at the time of decision. Height of the ‘Task’ and ‘Control’ bars corresponds to the average rating for these conditions. The figure also shows the individual subjects’ ratings for these, with the connecting line representing how the ratings varied between the two conditions. The error bar on the ‘Task – Control’ bar is the 95% confidence interval. B – Regions in mid-cingulum, middle frontal gyrus, and hippocampus showing significant correlation between the BOLD contrast, (Task – Control) contrast, and the free-will ratings, for Task – Control, in whole-brain exploratory correlational analysis. The plots accompanying the brain images are not independent results and are simply a visualization for the whole-brain correlation results shown in the images. For this reason, we do not report p-values for the correlations given the potential for selection bias (Vul et al. 2009).

**Figure 4.**
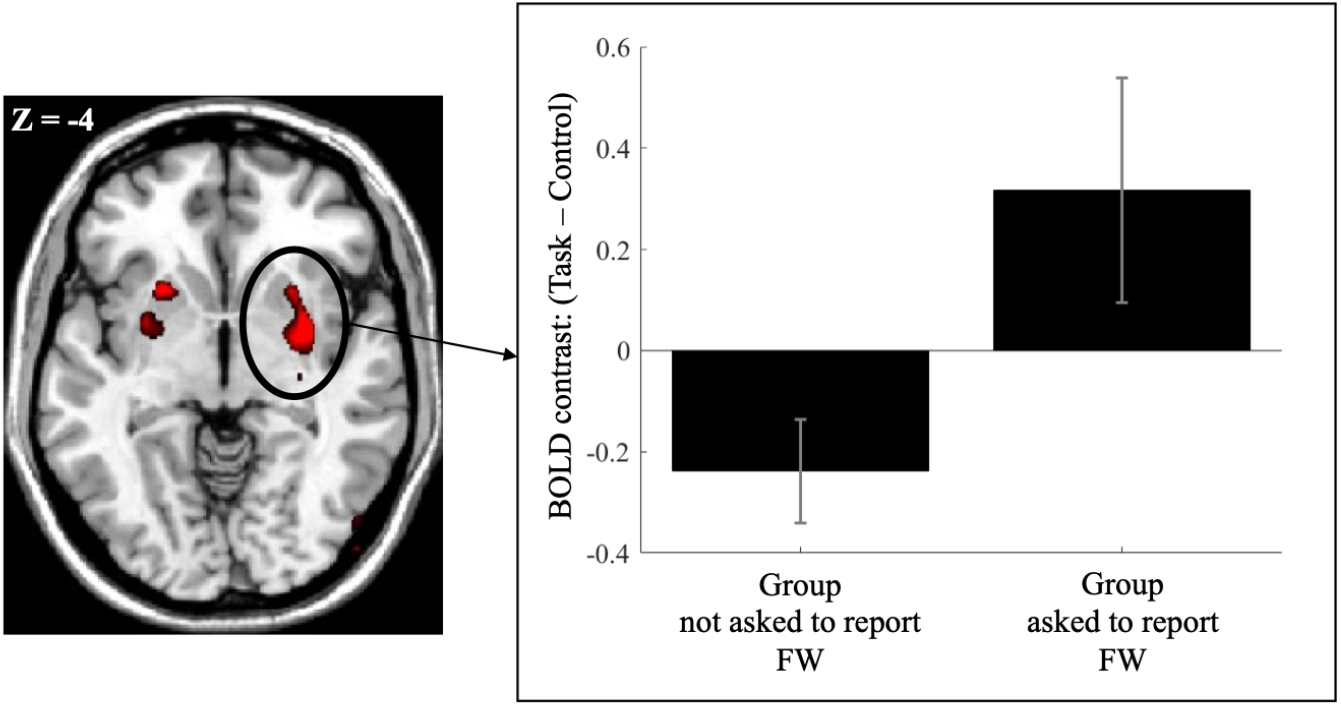
Cluster spanning putamen (striatum) and pallidum showing a significantly greater effect of (Task – Control) BOLD contrast in a two-sample exploratory whole-brain analysis for comparing the group of participants who were not asked to report free-will ratings and those who were asked to report the free-will rating. The figure on the left is not an independent result and simply represents the result on the right. The error bars depict confidence intervals from the mean contrast values across the group.

### Neural correlates of free-will ratings

Several regions showed correlations, across participants, between the difference in free will ratings for Task – Control and the BOLD response at the time of decision for Task – Control. Results of this whole brain exploratory analysis are shown in Figure 3 and Table 1.

**Table 1:** Clusters showing significant correlation in whole-brain exploratory correlation analysis of the (Task – Control) BOLD contrast at the time of decision and Task – Control self-reported free-will ratings. Cluster defining threshold (p-value) = 0.005. The p-value of clusters reported after FWE correction.

| Peak MNI coordinates |  |  |  |  |  |  |  |
| --- | --- | --- | --- | --- | --- | --- | --- |
| Region | Laterality | Cluster Size | X | Y | Z | Max stat t | P Cluster Corrected |
| Positive Correlation |  |  |  |  |  |  |  |
| Cingulate gyrus, BA 24, Mid-cingulum, BA 32 | Bilateral | 469 | -4 | 10 | 38 | 7.73 | 0.033 |
|  |  |  | 4 | -2 | 50 |  |  |
| Middle frontal gyrus (MFG) | Left | 639 | -26 | 18 | 36 | 6.04 | 0.006 |
| Negative Correlation |  |  |  |  |  |  |  |
| Hippocampus, Temporal pole, Parahippocampal gyrus | Left | 424 | -22 | -8 | -26 | 4.56 | 0.052 |

The ROI analysis of these clusters for the effect of (Task – Control) BOLD contrast revealed that the left MFG (mean contrast value = -0.96, p = 0.029, n = 12) and left hippocampus (mean contrast value = -0.62, p = 0.077, n = 12) clusters above show a significant and trending negative effect of the contrast, respectively. The significance was tested with a one-sample t-test of the average contrast values of these clusters for the participants. The cluster in the mid-cingulate overall shows a greater BOLD signal for task than the control but this is not significant for the sample size. We also looked at whether these clusters showed association with any of the behavioral measures and found that the cluster in hippocampus and parahippocampal gyrus showed a negative association with RT (r = -0.61, p = 0.036, n = 12).

Clusters showing significant positive and negative effects of the contrast (Task – Control) for the whole cohort (n = 30) are shown in supplementary tables S3 and S4. We considered each of these as a region of interest and assessed the correlation of the contrast values, (Task – Control), for the last twelve subjects who reported the free-will ratings with their free-ratings. Table 2 shows the clusters that showed significant linear relation (Pearson correlation) with the free-will ratings.

**Table 2:**
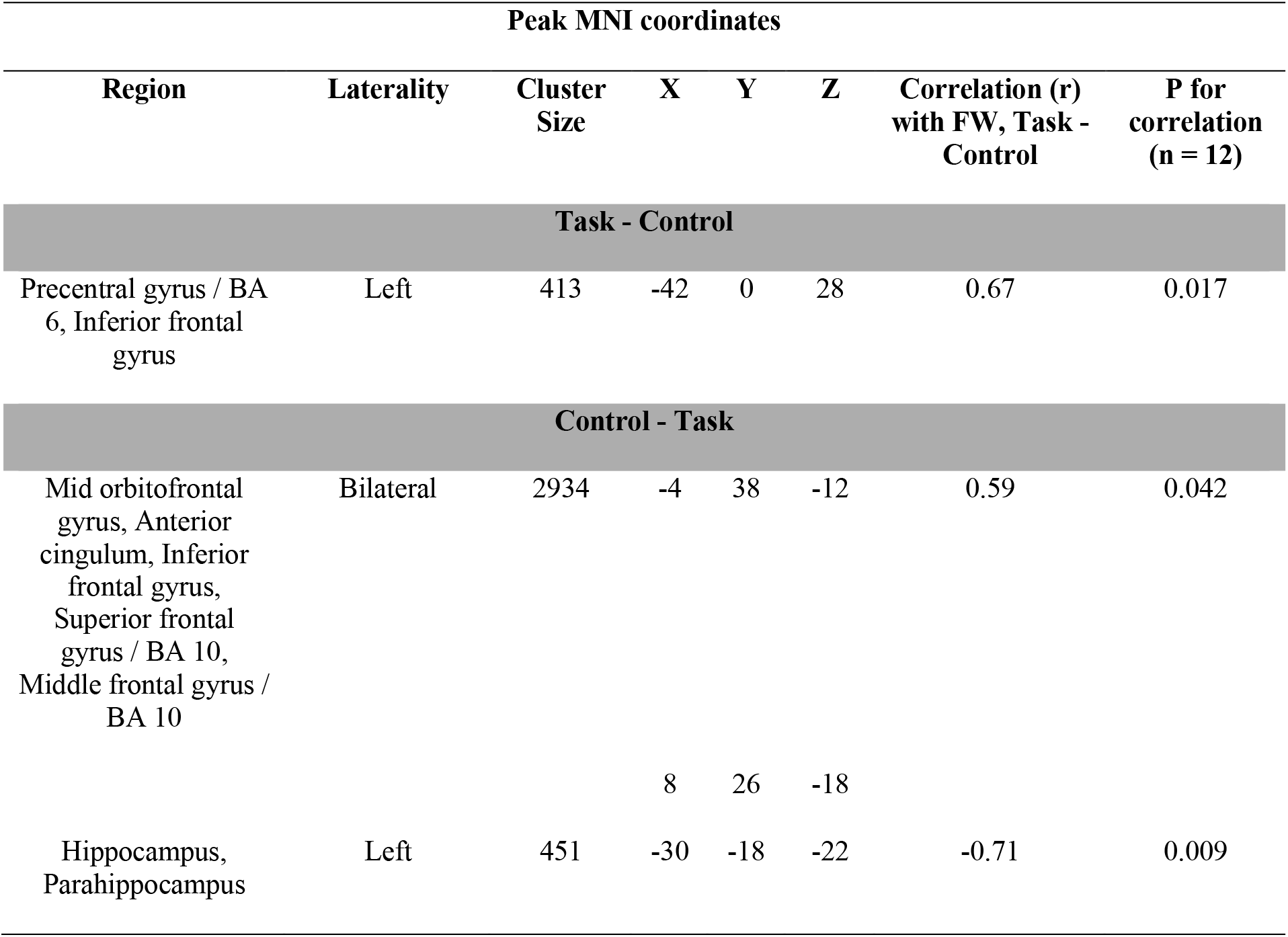
Clusters showing significant effect of the (Task – Control) or (Control – Task) in whole-brain GLM analysis with the entire cohort (n = 30) that also showed a significant association between the mean contrast values of (Task – Control) contrast with the free-will ratings (n = 12).

| Peak MNI coordinates |  |  |  |  |  |  |  |
| --- | --- | --- | --- | --- | --- | --- | --- |
| Region | Laterality | Cluster Size | X | Y | Z | Correlation (r) with FW, Task - Control | P for correlation ($n = 12$) |
| Task - Control |  |  |  |  |  |  |  |
| Precentral gyrus / BA 6, Inferior frontal gyrus | Left | 413 | -42 | 0 | 28 | 0.67 | 0.017 |
| Control - Task |  |  |  |  |  |  |  |
| Mid orbitofrontal gyrus, Anterior cingulum, Inferior frontal gyrus, Superior frontal gyrus / BA 10, Middle frontal gyrus / BA 10 | Bilateral | 2934 | -4 | 38 | -12 | 0.59 | 0.042 |
|  |  |  | 8 | 26 | -18 |  |  |
| Hippocampus, Parahippocampus | Left | 451 | -30 | -18 | -22 | -0.71 | 0.009 |

These clusters showing the main effect of the (Task – Control) or (Control – Task) BOLD contrast, were large in extent, so we also performed small volume correction of the whole-brain exploratory correlational analysis of BOLD contrast and free-will ratings within these clusters. This was done to localize any smaller regions within these clusters that may show a significant association with the free-will ratings. Please note that the effect of the contrasts was used to define the cluster which is independent of the effect that was tested with the small volume correction using these, which was the correlation between free-will ratings and the contrast values. We found that both the right and the left Insula to show significant positive correlation with the difference in free-will ratings after the small volume correction with the bigger Insula clusters that showed the main effect. This is shown in table 3.

**Table 3:**
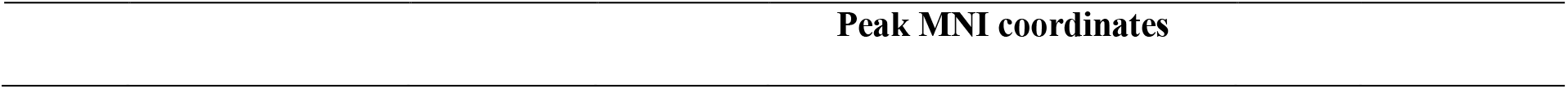
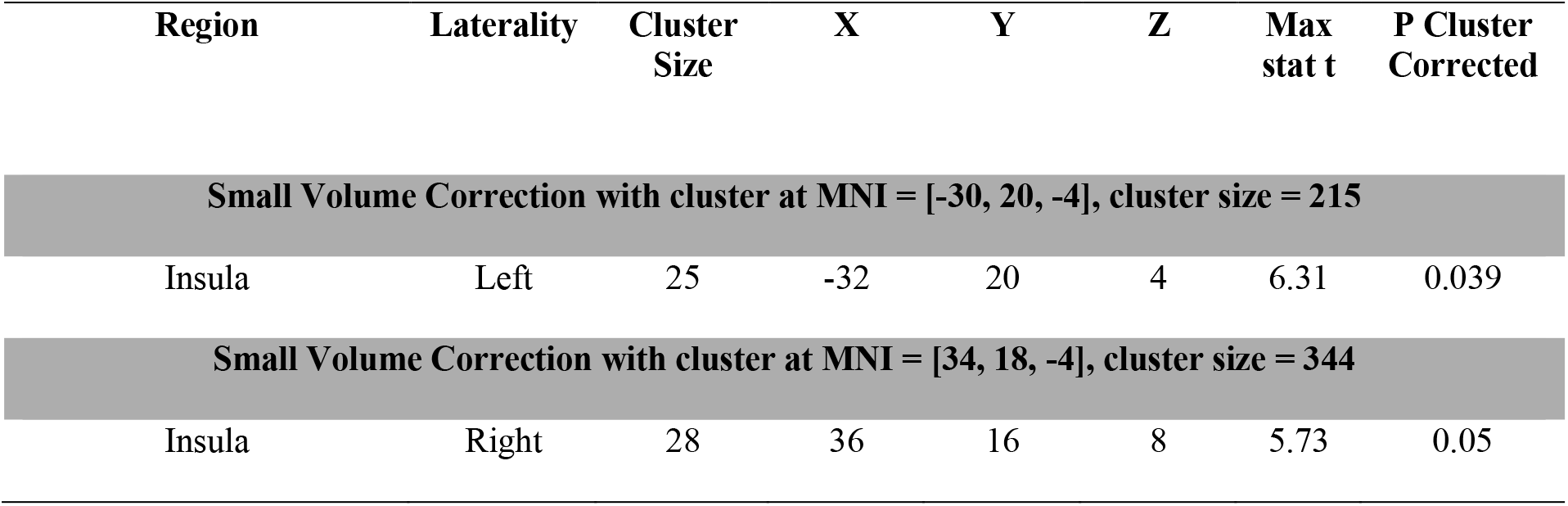
Clusters showing significant positive correlation, after small volume correction in whole-brain exploratory correlation analysis of the (Task – Control) BOLD contrast at the time of decision and Task – Control self-reported free-will ratings. Cluster defining threshold (p-value) = 0.005. The p-value of clusters is reported after FWE correction.

| Region | Laterality | Cluster Size | X | Y | Z | Max stat t | P Cluster Corrected |
| --- | --- | --- | --- | --- | --- | --- | --- |
| Small Volume Correction with cluster at MNI = [-30, 20, -4], cluster size = 215 |  |  |  |  |  |  |  |
| Insula | Left | 25 | -32 | 20 | 4 | 6.31 | 0.039 |
| Small Volume Correction with cluster at MNI = [34, 18, -4], cluster size = 344 |  |  |  |  |  |  |  |
| Insula | Right | 28 | 36 | 16 | 8 | 5.73 | 0.05 |

### Effects of monitoring free will experience

We next considered whether the requirement to report subjective experiences of free will could itself alter the neural processing. For this, we compared task-related BOLD signals between subjects who were asked to report free will experience vs. those who were not asked to report it. A whole-brain exploratory two-sample comparison for the (Task – Control) contrasts, with a cluster-defining threshold of p = 0.005, revealed a cluster in striatum that had a trending higher effect of the contrast (mean contrast value = 0.32, p = 0.009, n = 12) in participants who reported free-will while had a lower effect (mean contrast value = -0.24, p < 0.001, n = 18) in participants who were not asked to report free-will. This is shown in figure 2 and table 4 below.

**Table 4:**
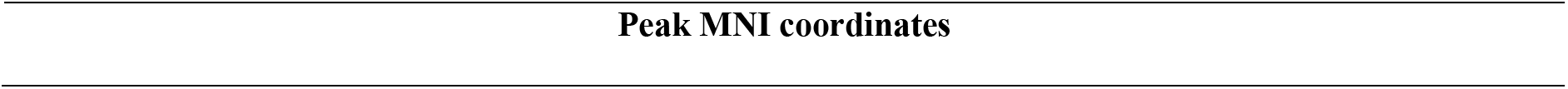
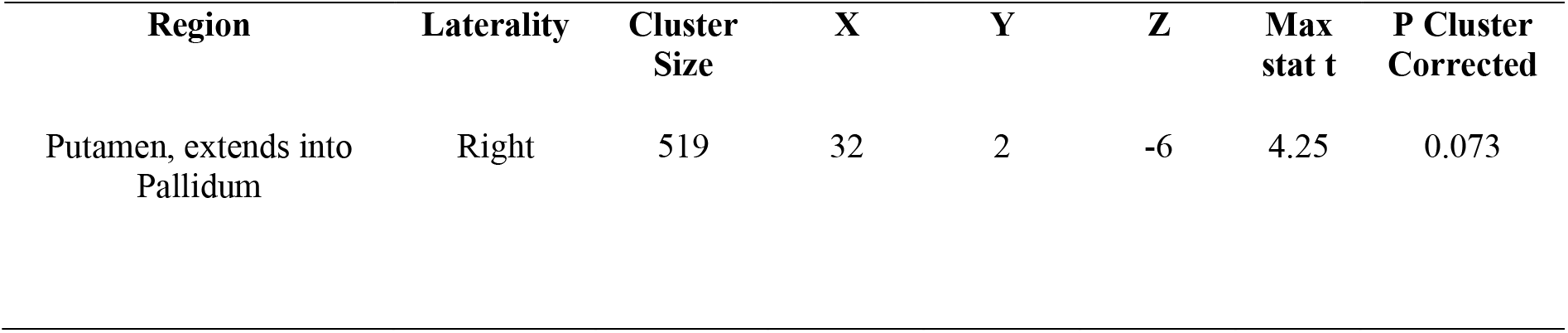
Cluster showing significant differences in the effect of (Task – Control) contrast at the time of decision, between the group that was asked to report free-will ratings (A) and one that was not asked to report this (B), such that A > B.

| Region | Laterality | Cluster Size | X | Y | Z | Max stat t | P Cluster Corrected |
| --- | --- | --- | --- | --- | --- | --- | --- |
| Putamen, extends into Pallidum | Right | 519 | 32 | 2 | -6 | 4.25 | 0.073 |

Among the clusters in table 1, a two-sample t-test of the contrast values for (Task - Control) BOLD contrast in the left MFG cluster showed a trend-like effect such that the contrast value for the group that reported free-will was lower or stronger negative (p = 0.097).

### Behavioral correlates of the free-will ratings

Free-will ratings, Task – Control, did not show any significant association with risk, conflict, consistency, or self-reported task difficulty, across participants. There was a positive trend between difference in free-will ratings and in RT (r = 0.46, p = 0.134), such that a greater RT for the Task compared to Control was associated with a greater free-will rating for the Task compared to Control.

## DISCUSSION

In this study, we set out to investigate neural and behavioral correlates of the experience of free-will. Healthy participants performed risky decision-making task where they chose between a risk-less sure option, that is ‘sure-thing’, and a gamble with two or three probabilistic outcomes (Figure 1C). In the task condition (Figure 1A), participants made their own choice and in the control condition (Figure 1C), they were instructed to choose a specific option. We collected their retrospective self-report ratings of experienced free-will during decisions at the end of each block (Figure 2). The sample size was small (n=12), so our findings should be considered preliminary.

We observed that the ratings for the subjective experience of free-will were reported to be different for different decision conditions, task and control. This indicates that this experience can vary for an individual and depends on the situation, comporting with the observations in previous literature (Filevich et al. 2013; Rens, Bode, and Cunnington 2018). Specifically, we found that the experienced free-will during the decisions in the Task condition was more than the Control condition. In both the conditions, participants saw two choice options, however, the task structure constrained the control condition such that only one option, the instructed choice, was viable for earning reward. This observation is in-line with Filevich et al. (2013) where they found that ratings for free-will correlated positively with ‘stimulus space’, which can be interpreted as a quantification of viable options in their study. This is also supported by Rens, Bode, and Cunnington (2018) where higher free-will ratings were associated with better representation of the choice alternatives.

These findings highlight a key aspect of experienced free-will, namely that simply being presented with different options does not necessarily lead to an experience of free-will and that the external and subjective constraints of the decision-environment need to be considered for identifying true or viable options. Our study formally shows and reiterates this ‘subjectivity’ of the experience of free-will as the variance observed in free-will ratings in our sample (Figure 3A). Across participants, the task instructions, number of options, and gamble stimuli were unchanged. Further, the sure-thing values were selected for each participant individually through a certainty equivalent task completed by the participants prior to the fMRI session with the intention to keep the decisions equally difficult for all participants during the fMRI session. Therefore, from the objective/experimenter perspective, across participants, the decision situations remained the same, and yet different individuals reported different levels of experienced free-will.

We also observed that one participant rated free-will to be higher for the control condition than the task condition. While this may merely be noise, this could also further highlight the subjectivity of the experience of free-will that is not necessarily dependent on the objective conditions. Such a rating may result from the participant being indifferent to the reward structure of the control task and not feeling constrained by the choice instruction, consequently experiencing a high level of free-will. In terms of behavior, the participant followed the choice instruction in the control task, but this does not necessarily negate the above interpretation as an experience of free-will may not necessarily translate to inconsistency in choices across multiple instances of the same decision scenario. Moreover, the task and control also differed in that the outcome of the choice could be predicted with certainty in the control condition while the outcome was uncertain in the task condition. Outcome uncertainty or controllability may additionally contribute to the experience of free-will and our further studies aim to investigate that.

We sought to also find behavioral markers of the experience of free-will. We defined free-will as the perceived capacity to choose freely among alternatives without constraints. Thus, one would predict that the free-will would positively be associated with choice conflict, or alternatively, negatively with consistency in task strategy, as several constraints (that is, low ‘free-will’) would lead to consistent choices or task-strategy. In our small sample, we did not find any significant or trend-like associations of free-will ratings with choice probabilities or risky behavior, conflict, and consistency in task strategy. In case of RT, we found a positive trend in its relationship with the ratings, such that participants who reported a higher experienced free-will also took longer to decide. It’s not surprising as previous studies (Proctor and Schneider 2018) have shown that a greater perception of alternatives is associated with a longer time to decide.

For the first-time to the best of our knowledge, here we also show the neural correlates of the subjective experience of free-will in economic decision-making. We found that a higher BOLD signal in bilateral mid-cingulate and left middle frontal gyrus (MFG), during value-based risky decisions (task), compared to control, was related to a greater experience of free-will. A greater BOLD activation in left hippocampus and parahippocampal gyrus was associated with a lower reported free-will. Firstly, we observed that not all of these regions showed the main effect of the (Task – Control) contrast. The region in hippocampus and parahippocampal gyrus showed only a trending effect of the main contrast while the region in mid-cingulate did not show a significant effect, in the ROI analysis. This shows that the neural correlates of the subjective experience of free-will do not necessarily coincide with the neural correlates of the free-will objectively defined, as the contrast between task and the control condition. This was also highlighted by Filevich et al. (2013). Secondly, among the regions showing the main effect of the (Task – Control) contrast, we show that the clusters in frontal cortex, anterior cingulate, and insula were associated positively with the subjective experience of free-will, while the clusters in hippocampus was associated negatively with it. Previous literature (Si, Rowe, and Zhang 2021; Thimm et al. 2012; Goldberg, Ullman, and Malach 2008; Welniarz et al. 2021) has implicated all of these regions in deliberative or intentional decisions, however, here we show how these clusters are associated with the subjective experience of free-will.

Further, while we have identified such regions, it remains to be investigated as to how this neural architecture contributes mechanistically to the experience of free-will. It is important to note that BOLD contrast, in the correlational analyses, corresponds to the time at which participants made the decisions while the reports of free-will were obtained afterwards, therefore, the regions identified above may not merely be involved in the process or act of reporting. Thus, subjective experience of free-will, reported by the participants, should be associated with the decision-making task itself, such as perceiving the availability of more than one task-strategies/options, or a perception of lower degree of internal/external constraints. Mid-cingulate or anterior mid-cingulate have been implicated in signaling response conflict (Carter et al. 1998) and its lower cortical thickness has been reported to be associated with higher self-reported helplessness (Salomons et al. 2012). It has also been associated with a greater involvement in decision conditions having external constraints to a lesser extent (Hoffstaedter et al. 2013; Modak and Brown 2026), and its stimulation associated with an experience of a ‘will to persevere’ (Parvizi et al. 2013). Thus, it appears that mid-cingulate may be associated with perceiving or identifying viable alternatives, which are more when constraints are low, which may explain why a greater BOLD signal in this cluster during decision-making may be associated with a greater self-reported free-will. Previous work has reported the BOLD signal in MFG to be associated with variance in outcomes and probability of loss (Fukunaga, Purcell, and Brown 2018; Modak et al. 2021), in high-level reasoning (Coricelli and Nagel 2009), in signaling option-availability (Rens, Bode, and Cunnington, 2018), and in choosing immediate rewards when left MFG specifically is disrupted (Figner et al. 2010). This suggests that the cluster in left MFG identified above, may be associated with the experience of free-will via representing viable options. A greater BOLD signal during decision-making in the cluster spanning hippocampus and parahippocampal gyrus associated with a lower level of subjective experience of free-will may be driven by its involvement in biasing the decision towards specific options (Li, Lu, and Zhong 2016; Wimmer and Büchel 2021; Wimmer and Shohamy 2012) and possibly reducing the extent to which other options are considered.

Moreover, since participants knew that they will be asked to report free-will ratings after the decisions, we also wondered whether the decision circuitry was altered from knowing that one will need to make this report later. To test this, we compared the effect of the BOLD contrast, (Task – Contrast) in the sample from whom these ratings were obtained with the one, from whom these ratings were not obtained. We identified a region in the striatum (putamen) extending into the pallidum, such that the effect of the (Task – Control) contrast was nearly significantly more positive in the group that reported free-will, compared to the group that was not asked to report it. It is also possible that merely being asked about free-will or knowing that one would need to report it, affected the experience of free-will itself – this possibility is hard to assess until there are ways to measure this experience without directly asking the participants. However, it does not seem that the region in the striatum/pallidum, identified as associated with the requirement to subsequently report free-will, was involved in altering the experience of free-will as this region did not show any association with the free-will ratings – such that there was no correlation between the ratings and the effect of the main contrast in this region. Therefore, it appears that this region might be involved in something that does not directly affect the experience of free-will, such as monitoring or pre-emptive introspection of this experience. This is in-line with previous neuroimaging work that has shown that striatum may be involved in signaling choice opportunity (Leotti, Cho, and Delgado 2015). Further, if striatum was monitoring the experience of free-will in our study, the BOLD signals for both the task and the control conditions must have been elevated (or reduced), in which case, there should not have been a change in (Task – Control) between the groups. However, our results are consistent with an interaction with the decision condition such that the monitoring or pre-emptive introspection of free-will may have elevated the BOLD signals in striatum/pallidum during the task condition much more than it did for the control condition.

## LIMITATIONS AND CONCLUSION

We only had two ratings of free-will for the task condition and one for the control condition per participant. This design choice was aimed at preventing the self-report of free-will from interfering with the task performance, but in future work, it may be better to get more ratings per participant to minimize the noise in self-report.

Further, our sample size is small, which comes with issues of alpha inflation. A replication with a larger cohort will be necessary before the specific results can be reliably generalized.

Nevertheless, this study provides a proof-of-concept that subjective reports of free will in value-based decisions relate meaningfully to BOLD activity. It further demonstrates that free-will can be examined at the level of experience and in the domain of non-motor decisions, even amid ongoing debate regarding its ontological status.

