## Supplementary Material for "Neural correlates of the subjective experience of free-will during value-based risky decisions: a pilot study"

1. Gambles

*Table S1: Details of the gamble options*

| Gambles | Values (V1 > V2 > V3) | | Probabilities | |
|---|---|---|---|---|
| **1** | V1 | 50 | P1 | 0.2 |
| | V2 | 15 | P2 | 0.5 |
| | V3 | -20 | P3 | 0.3 |
| **2** | V1 | 64 | P1 | 0.2 |
| | V2 | 10 | P2 | 0.6 |
| | V3 | -10 | P3 | 0.2 |
| **3** | V1 | 38 | P1 | 0.5 |
| | V2 | -7 | P2 | 0.5 |
| | V3 | NA | P3 | 0 |
| **4** | V1 | 40 | P1 | 0.5 |
| | V2 | 30 | P2 | 0.2 |
| | V3 | -10 | P3 | 0.3 |
| **5** | V1 | 38 | P1 | 0.3 |
| | V2 | 8 | P2 | 0.45 |
| | V3 | -1 | P3 | 0.25 |

2. Certainty equivalent task

The task is described below and also reported in our forthcoming paper (Modak and Brown, in prep).

We developed a custom webpage remote testing facility using a webpage using JavaScript, HTML, CSS, and PHP which was hosted on our lab's server to conduct a decision-making task, as described below, and a personalized link was sent to the participants where they could complete it. Participants completed a certainty equivalent task (Paulus and Frank 2006) and the sure thing values presented to them, and their choice responses were fitted using a logistic regression model for each of the five gambles. For a particular gamble, this model was used to find sure-thing values corresponding to choice probabilities, 0.35 and 0.65, referred to as $ST_{35}$ and $ST_{65}$.

The purpose of conducting this task was to obtain sure-thing values such that the actual sure-thing options were close to the certainty equivalents, so the decisions were not too easy for the participants.

During a task block, we randomly selected the gamble and then randomly selected one of the two sure-thing values corresponding to it with an added Gaussian noise ($\mu = 0$). The standard deviation was dependent on how steep the slope of the logistic model was around the $ST_{35}$ and $ST_{65}$ values such that it was equal to the average deviation in ST values required for a deviation of decision probability of 0.2 around $ST_{35}$ and $ST_{65}$. So, for a steeper slope, the standard deviation of the Gaussian used for adding the noise was smaller.

3. Payment

This is also reported in our forthcoming paper (Modak and Brown, in prep).

Participants received $10 for completing the certainty equivalent task (performed online, remotely) and $30/hour for the fMRI session. They also received a $10 bonus for arriving on time for the experiment.

Participants could also earn from the tasks, proportional to the points they earned during each of them:

- 0 to $5 for the certainty equivalent task
- 0 to 5$ for the decisions in task condition
- 0 to 2$ for the decisions in control condition

Payment was made in cash.

4. Free-will ratings

*Table S2: Free-will ratings provided by individual subjects for the task and control conditions*

| Subject # | Self-reported retrospective ratings of the level of free-will experienced during decision-making (Scale: 1-5; 1 – Not free at all; 5 – Completely free) | |
|---|---|---|
| | Task (Own choice) | Control (Instructed choice) |
| Sub 1 | 2 | 1 |
| Sub 2 | 1 | 1 |
| Sub 3 | 5 | 2 |
| Sub 4 | 4 | 2 |
| Sub 5 | 3.5 | 1 |
| Sub 6 | 3.5 | 3 |
| Sub 7 | 4.5 | 1 |
| Sub 8 | 2.5 | 2 |
| Sub 9 | 2 | 1 |
| Sub 10 | 3 | 1 |

| | | |
|---|---|---|
| Sub 11 | 4 | 4 |
| Sub 12 | 1.5 | 4 |

5. Supplementary fMRI result tables

The tables below display the clusters showing significant effects of the contrasts, Task – Control and Control – Task at the time of decision for the thirty participants, reprinted from our forthcoming paper (Modak and Brown, in prep).

<u>Task – Control</u>

*Table S3: Clusters showing the significant effect of the (Task – Control) contrast in a whole brain exploratory analysis*

| | | | | **Peak MNI coordinates** | | | |
|---|---|---|---|---|---|---|---|
| **Region** | **Laterality** | **Cluster Size** | **X** | **Y** | **Z** | **Max stat t** | **P Cluster Corrected** |
| Superior parietal lobule / BA 7, Precuneus / BA 7 / BA 39, Middle occipital gyrus | Bilateral | 8629 | 10 | -66 | 52 | 12.93 | <0.001 |
| | | | -10 | -68 | 56 | | |
| Thalamus, Medial global pallidus, Lentiform nucleus, Medial dorsal nucleus, Caudate, mid-brain, Bleeds into anterior cerebellum, Pallidum | Bilateral | 4026 | 10 | 6 | 0 | 8.28 | <0.001 |
| | | | -8 | 2 | -2 | | |
| Cingulate gyrus / Mid cingulum, BA 32, Middle frontal gyrus / BA 8, Supplementary motor area | Right | 1277 | 2 | 18 | 40 | 8.14 | <0.001 |
| Medial occipital gyrus / BA 18, Fusiform gyrus, Cerebellum_Crus1_L, | Left | 1731 | -44 | -58 | -18 | 7.54 | <0.001 |
| Insula, Inferior frontal gyrus / BA 47 | Left | 215 | -30 | 20 | -4 | 7.52 | 0.064 (FWE), 0.011 (FDR) |
| Precentral gyrus / BA 6, Inferior frontal gyrus | Left | 413 | -42 | 0 | 28 | 6.95 | 0.004 |
| Middle frontal gyrus / BA 6, Precentral gyrus | Right | 659 | 34 | 0 | 56 | 6.87 | <0.001 |

| | | | | | | | |
|---|---|---|---|---|---|---|---|
| Insula, Inferior frontal gyrus / BA 47 | Right | 344 | 34 | 18 | -4 | 6.77 | 0.01 |
| Cerebellum_9, Cerebellum, posterior lobe | Right | 173 | 10 | -60 | -50 | 6.61 | 0.123 (FWE), 0.02 (FDR) |
| Cerebelum_Crus1_L, Cerebellum posterior lobe, Vermis | Bilateral | 452 | 4 | -76 | -24 | 6.49 | 0.003 |
| | | | -6 | -74 | -28 | | |
| Middle frontal gyrus, Precentral gyrus | Left | 380 | -28 | -4 | 54 | 6.16 | 0.006 |
| Middle frontal gyrus / BA 46, Inferior frontal gyrus, | Right | 1236 | 48 | 10 | 24 | 5.85 | <0.001 |
| Cerebellum_9, Cerebellum anterior lobe, Vermis, Cerebellum posterior lobe | | 273 | 0 | -58 | -38 | 5.36 | 0.027 |
| | | | -12 | -52 | -44 | | |

<u>Control - Task</u>

*Table S4: Clusters showing the significant effect of the (Task – Control) contrast in a whole brain exploratory analysis*

| | | | **Peak MNI coordinates** | | | | |
|---|---|---|---|---|---|---|---|
| **Region** | **Laterality** | **Cluster Size** | **X** | **Y** | **Z** | **Max stat t** | **P Cluster Corrected** |
| Cingulate gyrus / BA 31, Mid cingulum, Precuneus, Superior temporal gyrus / BA 39, Middle temporal gyrus (Bleeds into Inferior and superior temporal gyrus / BA 21), Supramarginal gyrus / BA 40, Inferior parietal lobule, | Bilateral | 19385 | 64 | -38 | 30 | 8.21 | <0.001 |
| | | | -52 | -8 | -26 | | |
| Mid orbitofrontal gyrus, Anterior cingulum, Inferior frontal gyrus, Superior frontal gyrus / BA 10, Middle frontal gyrus / BA 10 | Bilateral | 2934 | -4 | 38 | -12 | 6.11 | <0.001 |
| | | | 8 | 26 | -18 | | |

| | | | | | | | |
|---|---|---|---|---|---|---|---|
| Hippocampus, Parahippocampus | Left | 451 | -30 | -18 | -22 | 5.46 | 0.003 |
| Superior occipital lobe, Cuneus, BA 18 | Right | 544 | 14 | -82 | 26 | 5.41 | 0.001 |
| Superior occipital lobe, Cuneus, BA 18, BA 19 | Left | 253 | -12 | -90 | 22 | 5.07 | 0.036 |
| Sub-gyral, Caudate | Right | 161 | 24 | 8 | 20 | 4.10 | 0.149 (FWE), 0.056 (FDR) |

### References

Modak, Priyamvada, and Joshua W. Brown. 2021. “Motivationally objective versus subjective decision-making: Neural correlates, behavior, and self-report.”*In Preparation.*
